# Novel bile salt analogs reduce lipid accumulation in liver cells with potential to treat both metabolic dysfunction-associated steatotic liver disease and Clostridioides difficile infection

**DOI:** 10.64898/2026.06.11.731657

**Authors:** Defeng Cai, Huy Nguyen, Yang Zhang, Shiv Sharma, Angel Schilke, Rola Raychouni, Efren Heredia, Ernesto Abel-Santos, Steven Firestine, Wanqing Liu

## Abstract

Metabolic dysfunction-associated steatotic liver disease (MASLD) and *Clostridioides difficile* (*C. difficile*) infection (CDI) are clinically associated, yet there is limited effective treatment for both diseases. Bile salt analogs (BSAs) have demonstrated potential in treating either MASLD or CDI. We screened a library of BSAs (n=112) previously synthesized as potential inhibitors of *C. difficile* spore germination, for their therapeutic potential in reducing intracellular accumulation of fatty acids in HepG2 cells as candidates for prevention and treatment of both MASLD and CDI. The screening was based on an *in vitro* model established by incubating HepG2 cells with free fatty acids, with obeticholic acid (OCA), a known BSA with anti-MASLD activity as a control. Gene and protein expressions were quantified to validate the treatment effect. We found that compounds C13, C24, C25, C74, C98, and C101 demonstrated significant effectiveness in both preventing the intracellular accumulation of lipids and removing pre-loaded cellular lipids. Gene expression analysis showed that C24, C25, and C74 produced a similar pattern characterized by a robust induction of FGF21 expression, while C13, C98, and C101 produced a transcription pattern that mirrors the effect of OCA. Structurally, while C13, C24, and C25 do not display drug-like properties, C74, C98, and C101 are drug-like and share a similar structure. Interestingly, C101 is a potent inhibitor of *C. difficile* spore germination. OCA shows a weak anti-gemination effect. Our study identified lead compound candidates for the development of novel therapeutics capable of treating both MASLD and CDI.

**Significance statement:** The clinical association between MASLD and CDI remains an unmet need for dual acting therapeutic strategies. Given the reported potential of BSA, we screened 112 previously synthesized as potential inhibitors of *C. difficile* spore germination, for their therapeutic potential in reducing intracellular accumulation of fatty acids in HepG2 cells. Our study identified compounds that effectively reduce intracellular lipid accumulation and inhibit *C. difficile* spore germination. These results nominate lead candidates for developing dual-acting therapeutics targeting both MASLD and CDI.

## INTRODUCTION

Metabolic-dysfunction associated steatotic liver disease (MASLD) is the most common chronic liver disorder globally. ^1^ MASLD is characterized by intracellular fat accumulation as hepatosteatosis in individuals with metabolic dysfunction. ^2^ Without proper intervention, hepatosteatosis can progress to a more severe form of liver disorder called metabolic-associated steatohepatitis (MASH) which is characterized by histological changes including steatosis, lobular inflammation, and fibrosis. If left untreated, MASH can further advance to cirrhosis and even liver cancer. ^2 3^

The detailed molecular mechanisms underlying the development of the histological spectrum of MASLD are not completely understood. However, it is believed that liver steatosis (the accumulation of neutral lipids inside hepatocytes) is the initial step for the development of MASH. ^3 4^ Therefore, targeting lipid accumulation in hepatocytes is deemed as a fundamental strategy to prevent MASLD from progressing and for the treatment of MASH.

Despite the rising prevalence of MASLD and MASH in recent decades, there are few approved pharmacological treatments. Resmetirom, a thyroid hormone beta receptor (THR-β) agonist, has become the first FDA-approved drug for treating MASH. ^5^ Mechanistically, Resmetirom was found to increase autophagy, mitophagy, mitochondrial biogenesis, and lipids β-oxidation in liver cells. ^6 7^ The GLP-1 receptor agonist, semaglutide was also approved for treating MASH. However, GLP-1 receptor expression in the liver and hepatocytes is absent or extremely low. ^8–15^ The impact of semaglutide on improving liver histology was primarily deemed as an indirect effect of weight loss or alteration of metabolic signaling. ^16–18^

Numerous candidate drugs have been tested in pre-clinical and clinical studies for MASH treatment. These interventions target different pathogenic pathways involved in MASLD and MASH. For example, the antioxidant properties of vitamin E have demonstrated clinical benefit for MASH treatment via normalization of increased oxidative stress. ^19^

Recently it was shown that administration of fibroblast growth factor 21 (FGF21) reverses diet-induced MASH in rodents through coordinated actions on the CNS and liver ^20^ Furthermore, FGF21 analogs modulate multiple metabolic pathways among multiple target tissues (e.g., liver and adipose tissues). These compounds exert metabolism modulation properties by binding to the FGFR1c and β-Klotho receptor complex. ^21 22^ These results in the balancing of glucose and lipids metabolism, improving insulin sensitivity and energy expenditure, as well as ameliorating inflammatory and fibrotic responses. ^21 23^ Although no FGF21 analogs have been approved for clinical use thus far, two drugs (efruxifermin and pegozafermin) are currently in clinical trials for patients with MASH. ^24^

*Clostridioides difficile* infection (CDI) is responsible for up to 25% of all antibiotic-associated diarrheas. ^25^ CDI management is complicated by the ability of *C. difficile* to form dormant and resistant spores. *C. difficile* spores persisting in hospitals result in cross-infections. ^26^ Similarly, *C. difficile* spores remaining inside patients’ intestines can lead to relapse. ^27^ *C. difficile* spores do not cause infection but can germinate in the presence of bile salts into toxin-producing vegetative cells. ^28 29^ Bile salts are synthesized in the liver and secreted into intestine to facilitate dietary lipids absorption. It has been well-established that gut microbiota and the liver metabolism dynamically interact, during which bile salt homeostasis is one of the key mediators via the enterohepatic circulation (EHC). ^30 31^ Altered bile salt profiles are directly associated with MASLD/MASH, ^32 33^ as well as CDI. ^34^

Bile acids are biosynthesized in a series of cytochrome P450-mediated reactions (e.g. CYP7A1, CYP27A1, and CYP8B1). ^32 35^ The farnesoid X receptor (FXR) is a critical hepatic regulating transcription factor which, once activated by bile acids, transduces feedback signals to inhibit the expression of CYP7A1 and regulate bile acid homeostasis. ^32 35^ The majority of secreted bile acids from the liver are re-absorbed back to the liver from the ileum via the EHC. ^32 35^

Bile salts also play an important role in homeostasis and metabolism of both fatty acids and cholesterol in the liver. ^32 33 35 36^ Similarly, bile salts have a reciprocal interaction with gut microbiota: while bile acids can modulate the microbiome profile, gut bacteria are involved in the metabolism of bile acids. Intriguingly, the gut microbiome profile is also a crucial factor that alters susceptibility to MASLD. ^37^

The bile salt analog (BSA) obeticholic acid (OCA), a potent FXR agonist, has demonstrated excellent potential for treating MASLD and MASH, though certain side effects (*e.g.*, pruritus) remain as a concern. ^38 39^ Mechanistically, OCA was found to modulate FXR activation in the liver, which reduces hepatic lipogenesis, gluconeogenesis, inflammation, and fibrotic reactions. In parallel, FXR activation by OCA in the intestine induces the production of FGF19, an important metabolic modulating hormone, balancing bile acid homeostasis. ^40 41^ Hence, bile acids and their synthetic analogs are attractive drug candidates for treating MASLD and MASH. ^42^ Following the example of OCA, the use of synthetic bile acid derivatives for lipid regulation to treat MASLD continues as an active research field.

Many of the BSAs that activate FXR are derived from the naturally occurring chenodeoxycholic acid (CDCA). Interestingly, CDCA is a natural inhibitor of *C. difficile* spore germination. ^43 44^ Furthermore, OCA has shown beneficial impact on an obese model of murine CDI. ^45^

Since bile salt-mediated *C. difficile* spore germination is a necessary step in CDI onset, we previously developed libraries of BSAs that could block *C. difficile* spore germination and thus, prevent CDI ^46^. Indeed, we found a series of synthetic BSA anti-germinants that also prevent CDI in rodent models. ^29 47 48^

Recent epidemiological studies demonstrate that MASLD/MASH and CDI (or its clinical outcomes) are strongly associated. ^49–52^ However, the causal relationship between MASLD and CDI remains largely elusive. It is also unclear whether treatment for one disease would prevent another.

Given the potential structural overlap between *C. difficile* spore germination inhibitors and FXR agonists, we elected to screen the library of synthetic BSAs (n=112) with the goal to identify novel drugs that could prevent and/or reduce cellular hepatosteatosis as potential therapy for MASLD and MASH. Our primary goal of this screening is to examine the lipid-reducing capacity of these compounds in the HepG2 cell model. Our study found some promising candidate compounds that can reduce intracellular lipid accumulation possibly by activating either FGF21 or FXR signaling. Interesting, some of these compounds have demonstrated strong activity in inhibiting CDI suggesting that there could be multi-purpose BSAs that would be useful in treating multiple diseases.

## MATERIALS AND METHODS

### Materials

The HepG2 cell line was originally obtained from American Type Culture Collection (Manassas, VA, USA). Bile salt analogues were synthesized in the Firestine lab at Wayne State University according to previous methods. ^46^ All compounds were dissolved in dimethyl sulfoxide (DMSO) to prepare 5 mM stock solutions. Cell media (Dulbecco Modified Eagle’s medium, DMEM), bovine serum albumin (BSA), Triton X-100, DMSO, GlutaMax, Ciprofloxacin, fetal bovine serum (FBS), TRIzol Plus RNA purification kit, High-Capacity cDNA Reverse Transcription kit, RIPA buffer, protease inhibitor cocktail, sodium dodecyl sulphate-polyacrylamide gradient gels (4%-12%), and the BCA protein quantification kit, pre-stained protein ladders, nitrocellulose membranes, non-fat milk, TBST buffer, and the ChemiGlow West Chemiluminescence Substrate kit for Western blotting were purchased from Thermo Fisher Scientific (MA, USA). SYBR green Master Mix real-time PCR kit was purchased from Bio-Rad (CA, USA). The LabAssay™ Triglyceride assay kit was purchased from FUJIFILM Wako Pure Chemical Corporation (VA, USA). Palmitic acid (PA), oleic acid (OA), BODIPY, Hoechst 33342, and silica thin layer chromatography (TLC) assay kit was purchased from MilliporeSigma (MO, USA). The primary antibodies for detecting FGF21 and GAPDH were obtained from Abcam (MA, USA) (ab171941) and Cell Signaling (MA, USA) (2118s), respectively. The secondary antibody was also purchased from Cell Signaling (MA, USA) (7074s).

### HepG2 cell culture

The human hepatoma cell line HepG2 was maintained in Dulbecco Modified Eagle’s medium (DMEM) with 1.0 g/L D-glucose and 2mM L-glutamine. The medium was supplemented with 10% fetal bovine serum, 1% GlutaMax, and Ciprofloxacin (10µg/ml). Cells were maintained in T75 cell culture flasks (Thermo Fisher Scientific, CA, USA) in 10 mL of medium in an incubator at 37 °C with a humidified atmosphere and 5% CO_2_. Upon reaching ∼70% confluency, cells were sub-cultured by washing twice with 10 mL of phosphate buffered saline (PBS) with 5 mM EDTA for 2 min at room temperature. Cells were then detached from the flask by repetitive aspirating with a pipette. Suspension cells were then centrifuged at 500 × g for 5 minutes. After decanting, the cell pellet was resuspended in growth medium. Cell viability of each passage was determined by trypan blue staining using a hemocytometer.

### Preparation of palmitic acid (PA) and oleic acid (OA) solutions

PA and OA were dissolved in deionized H_2_O at 60 . The resulting solutions were further solubilized in PBS containing 10% fat-free BSA to make 20 mM stock lipid solutions. Solutions were filtered with a syringe filter (Thermo Fisher Scientific, MA, USA) and diluted to the corresponding lipid concentrations as required.

### Establishment of the high-throughput lipid analysis cell model using BODIPY/Hoechst staining

Lipid accumulation in HepG2 cells was induced by treating the cells with mixed PA and OA, as previously described ^53^. Briefly, cells were seeded at 3,000 cells/well in 96-well plates containing 200 µl of growth media and grown to approximately 70% confluency. Cell cultures were then treated with mixtures containing either 100 µM PA + 200 µM OA, 200 µM PA + 400 µM OA, 300 µM + PA 600 µM OA, or 400 µM PA + 800 µM OA. Cells were incubated for 24 hrs with their corresponding lipid mixtures. Cells were then washed twice with PBS. Washed cells were treated with a solution containing 5 μg/ml BODIPY (to stain and measure total intracellular neutral lipids) and 1 μg/ml Hoechst 33342 (to stain and measure the number of cell nuclei) and incubated for 30 min at 37 . Stained cells were washed three times with PBS. Fluorescence intensity for BODIPY and Hoechst 33342 was quantified by using a CLARIOstar plate reader (BMG Labtech, NC, USA) at 477 nm and 355 nm, respectively. The BODIPY/Hoechst fluorescence ratio of each well was calculated to normalize for intracellular lipid accumulation. A standard curve between the treatment concentration and the fluorescence ratio was determined to examine detection linearity and range between fluorescence ratios and intracellular levels of accumulated lipids.

### Triglyceride (TG) level measurement in cells and culture media

The level of accumulated lipids was further validated by quantifying the total triglycerides (TG) level in each well, normalized to the total protein level in each well. The TG content in HepG2 cells was measured in a microplate following manufacturer’s instructions. Briefly, after treatment, cells were seeded at 3000 cells/well and grown until ∼70% confluency. Cells were then washed twice with PBS, followed by lysis on ice with 1% Triton X-100 in PBS supplemented with protease inhibitors (Thermo Fisher Scientific, MA, USA). Supernatants were collected after centrifuging cell lysates at 12,000 g for 10 min. The TG levels were then measured colorimetrically by mixing 2µl of cell lysate and 300µl of Chromogen buffer provided in the kit and followed by an incubation at 37 for 5 min. Absorbance from cell lysates and standard samples was then read in the CLARIOstar plate reader (BMG Labtech, NC, USA) at 570 nm. TG levels were calculated based on interpolation into the standard curve.

To normalize TG concentrations, total protein was measured using the same cell lysate obtained above. Briefly, protein concentration of the supernatant of each sample was determined using the BCA protein assay kit based on a standard curve by following the manufacturer’s instructions. Total TG concentration was normalized by dividing the TG concentration by total protein concentrations obtained from matched cell culture lysates.

### Screening bile salt analogs for preventing lipids accumulation

To screen for lipid-accumulation prevention, HepG2 cells were grown in 96-well plates, as above. Cells with 70% confluence were supplemented with either DMSO (negative accumulation control) or 100 µM PA + 200 µM OA in DMSO (positive accumulation control). The compound-treatment wells were immediately treated with either 0 (*i.e.,* the positive accumulation control noted above), 6.25, 12.5, 25, 50, or 100 µM of a candidate compound in DMSO. Cells were grown 24 hrs in supplemented media to allow for intracellular lipid accumulation. After incubation, intracellular lipid accumulation was determined as above.

### Screening of active bile salt analogs for lipid removal

To screen for lipid-removal activity, HepG2 cells were grown in 96-well plates, as above. Cells with 70% confluence were supplemented with either DMSO (negative accumulation control) or 100 µM PA + 200 µM OA in DMSO (24 hrs. accumulation control), as above. After 24 hrs incubation, cells were then washed with PBS to remove extracellular PA and OA. Washed cells were maintained in regular media and treated with either 0 (*i.e.,* the 24 hrs accumulation control noted above), 6.25, or 12.5 µM of an active compound in DMSO for an additional 24 hrs. As a second control, separate wells were supplemented with 100 µM PA and 200 µM OA for the entire duration of the experiment (48 hrs accumulation control). After incubation, intracellular lipid accumulation was determined as above.

### Dose-dependent effect of active bile salt analogs on lipid removal

To quantify the potency of these compounds in reducing pre-loaded lipids, HepG2 cells were treated as above. After 24 hrs treatment, washed cells were then maintained in a regular media and treated with either 0 (*i.e.*, the 24 hrs accumulation control), 2, 4, 8, 16, and 32 µM of an active compound for an additional 24 hrs. Fluorescence ratio of BODIPY/Hoechst was then quantified for cells treated with each compound concentration. The concentration of each compound required to reduce lipid accumulation to 50% of the 24 hrs accumulation control (EC_50_) were calculated using the “Dose-Response – Inhibition” function under Nonlinear Regression of GraphPad Prism 8.0 (GraphPad Software, MA, USA). Data was also plotted by using GraphPad Prism 8.0.

### Time-dependent effect of active bile salt analogs on lipid removal

The same model for lipids removal as detailed above was used for testing the time-dependent lipid-lowering effect of selected candidate compounds (C13, C24, C25, C74, C98, and C101). The entire test was performed for a duration of 48 hrs. Briefly, HepG2 cells with treated with DMSO (negative accumulation control) or with 100 µM PA + 200 µM OA in DMSO for 24 hrs. (24 hrs accumulation control). After 24 hrs of lipids accumulation, the control groups were treated with a second aliquot of neat DMSO (control group) or with 6.25 µM of an individual compound in DMSO. After 4, 8, 12, or 24 hrs of treatment, intracellular lipids content was measured based on the fluorescence ratio, as above.

### Fluorescence microscopy for lipid droplets formation

HepG2 cells were cultured following the same procedures as the lipid removal experiments. After treatment, cells were stained with BODIPY and Hoechst as above. Fluorescence signals were visualized with a ZEISS laser scanning microscope 800 confocal microscopy (Carl Zeiss, Germany) to compare the lipid droplets formation between groups. Image J software ^54^ was used to quantitatively analyze accumulated neutral lipids in each group.

### Thin-layer chromatography (TLC) for lipidomic analysis

HepG2 cells were cultured following the same procedures as the lipid removal experiment but using a single 6.25µM concentration of selected compounds. OCA (6.25 µM) was used as a positive control. After treatment, cell pellets were collected, and total lipids were extracted using a previously published protocol ^55^. Briefly, after washing cells with PBS, cells were digested with trypsin and re-suspended in PBS. A standardized number of 5×10 cells was used for lipid extraction. Methanol, water, and chloroform were sequentially added to the cell pellet, followed by vortexing to resuspend the cellular mixture. The cell suspension was centrifuged at 3000 g for 5 minutes. After centrifugation, the lower chloroform-rich phase that contains neutral lipids was collected and further washed with pure methanol followed by water to remove impurities. The chloroform phase was centrifuged again and separated from polar solvents. The chloroform phase was then air-dried in a chemical hood at room temperature.

For TLC, the dried lipid extract was resuspended in 150 µL chloroform. A 30 µl aliquot of each lipid solution was spotted onto a silica gel TLC plate 2 cm from the bottom. Plates were developed with a hexane:diethyl ether:acetic acid (80:20:2) mixture until the solvent front reached approximately 1.5 cm from the top of the plate (∼12 min). The chromatographic plate was then air-dried for 10 minutes. The TLC plates were immersed in a staining solution (0.005% Primulin dissolved in 80:20 acetone/water) for 10 seconds. The plate was then air-dried, visualized under UV light (302 nm) and imaged. Lipids were quantified using the Image J software. ^54^

### Gene expression analysis

To determine the effect of lipid-lowering compounds on HepG2 cell transcriptome, cells in compound-treatment groups were co-treated with PA (100µM) + OA (200µM) and an individual compound at a fix 6.25µM concentration. OCA (6.25 µM) was used as treatment control. negative accumulation control cells were treated with neat DMSO. positive accumulation control cells were treated with PA (100µM) + OA (200µM) in DMSO. After 24 hrs, cells were collected and total RNA was extracted using Trizol, following the manufacture’s guidelines. Total RNA was then reverse-transcribed following manufacturer’s instructions. Quantitative PCR (qPCR) was used to quantify the expression level of twenty-six HepG2 genes related to various pathways of lipid metabolism. The *GAPDH* gene was used as an internal qPCR reference. All primers were selected from Primerbank ^56^, with sequences shown in **Table S1**. Results of all qPCR runs were analyzed using the double delta Ct (2^−ΔΔCt^) method. Gene expression comparison, heatmaps and clustering analyses were conducted with the R package (https://www.R-project.org/).

### Western blot analysis

Western blot assays for FGF21 and GAPDH were performed based on previously established methods ^53 57^. Briefly, cells were treated following the same procedures as for the gene expression experiment, described above. Total cellular protein was obtained as described above. A 20 μg protein aliquot of each sample along with a pre-stained protein ladder were separated by a 4%-12% SDS-PAGE. Proteins were then transferred to a nitrocellulose membrane and blocked with 5% non-fat milk in TBST for overnight at 4°C. FGF21 and GAPDH were respectively blotted with their primary antibodies overnight at 4°C. Following washing with TBST, the membrane was further blotted with the secondary antibody for 1 hr at room temperature. Protein bands were then detected with the ChemiGlow West Chemiluminescence Substrate kit and imaged using the Bio-Rad ChemiDoc Imaging System (Bio-Rad, MI, USA).

### Anti-germination activity of OCA

*C. difficile* strain R20291 was a kind gift of Prof. Nigel Minton (University of Nottingham). *C. difficile* spores were prepared according to previously published protocols ^46^. Purified spores were prepared for germination according to previously published protocols ^46^. Briefly, spores were diluted to an optical density at 580 nm (OD_580_) of 1.0 with 100 mM sodium phosphate buffer, pH 6.0, supplemented with 5 mg/mL sodium bicarbonate. Increasing concentrations of OCA were added, in triplicate, to a 96-well plate. To each well, 6 mM taurocholate and 12 mM glycine were added, followed by 180 μL of spores for a final volume of 200 μL. The extent of *C. difficile* spore germination was assessed as the decrease in OD_580_ over time. Thus, OD_580_ was measured once every minute for 2 hrs and normalized using the OD_580_ obtained at time zero [relative OD_580_ = OD_580_(t)/OD_580_(t_0_)].

The resulting germination curves were used to determine percent germination by comparing the slope from the linear part of the early germination curves of each OCA concentration to the slope from the germination curve in the absence of OCA. Percent germination rates were then plotted against their corresponding OCA concentration. The resulting sigmoidal curve was analyzed in SigmaPlot version 14 (Grafiti LLC., NY, USA) by fitting with the four-parameter logistic function to obtain the IC_50_ value for OCA. ^48^

### Assessment of drug-like property using SwissADME

The chemical structure for C13, C24, C25, C74, C98 and C101 were converted to SMILES. The SMILES representation for all compounds was copied into the appropriate location in SwissADME (swissadme.ch). ^58^ A series of calculated and predicted properties for each molecule was generated including lipophilicity and 5 predictors of drug likeness. Any prediction which listed a violation of at least one of their parameters was taken as an indication that the compound was not drug-like.

### Statistical analysis

Statistical differences between treatment groups were first examined with single-factor One-way ANOVA. ANOVA results with p<0.05 were further analyzed *post hoc* using the Tukey test for pairwise comparison between groups using the GraphPad Prism 8.0 (GraphPad Software, MA, USA). The corrected p<0.05 based on the Tukey test was used as the cut-off for statistical significance. P-values were only reported for selected groups that are relevant to the effect of treatment.

## RESULTS

### Establishment of the lipid analysis model

We first optimized a lipids quantification method by using the BODIPY/Hoechst fluorescence ratios. using different PA/OA concentration revealed that the BODIPY/Hoechst ratio reflects the dose-dependent accumulation of intracellular lipids in HepG2 cells (r^2^=0.97 p<0.0001, **Figure S1A**). To further validate this model, the TG: total protein ratio in each well was also measured, which confirmed that supplementation of HepG2 cells with PA/OA led to intracellular lipid accumulation (r^2^=0.96, p<0.0001, **Figure S1B**). All subsequent lipid accumulation experiments were quantified using BODIPY/Hoechst ratios.

### Screening of 112 compounds in preventing lipids accumulation in HepG2 cells

Previously, a library of bile salt analogs was prepared in the Firestine laboratory to identify inhibitors of *C. difficile* spore germination. ^46^ Since earlier studies have shown that bile salt analogs can regulate lipid homeostasis, we decided to investigate this library (N=112) of bile salt analogs for anti-steatosis activity. Of the 112 compounds tested, six analogs (C13, C24, C25, C74, C98 and C101) showed a significant dose-dependent effect for preventing intracellular lipid accumulation in HepG2 cells (**Figure 1)**. Chemical structure of these compounds is included in **Figure S2**. None of the six active compounds showed significant cytotoxicity, even at the highest concentration tested (data not shown).

**Figure 1.**
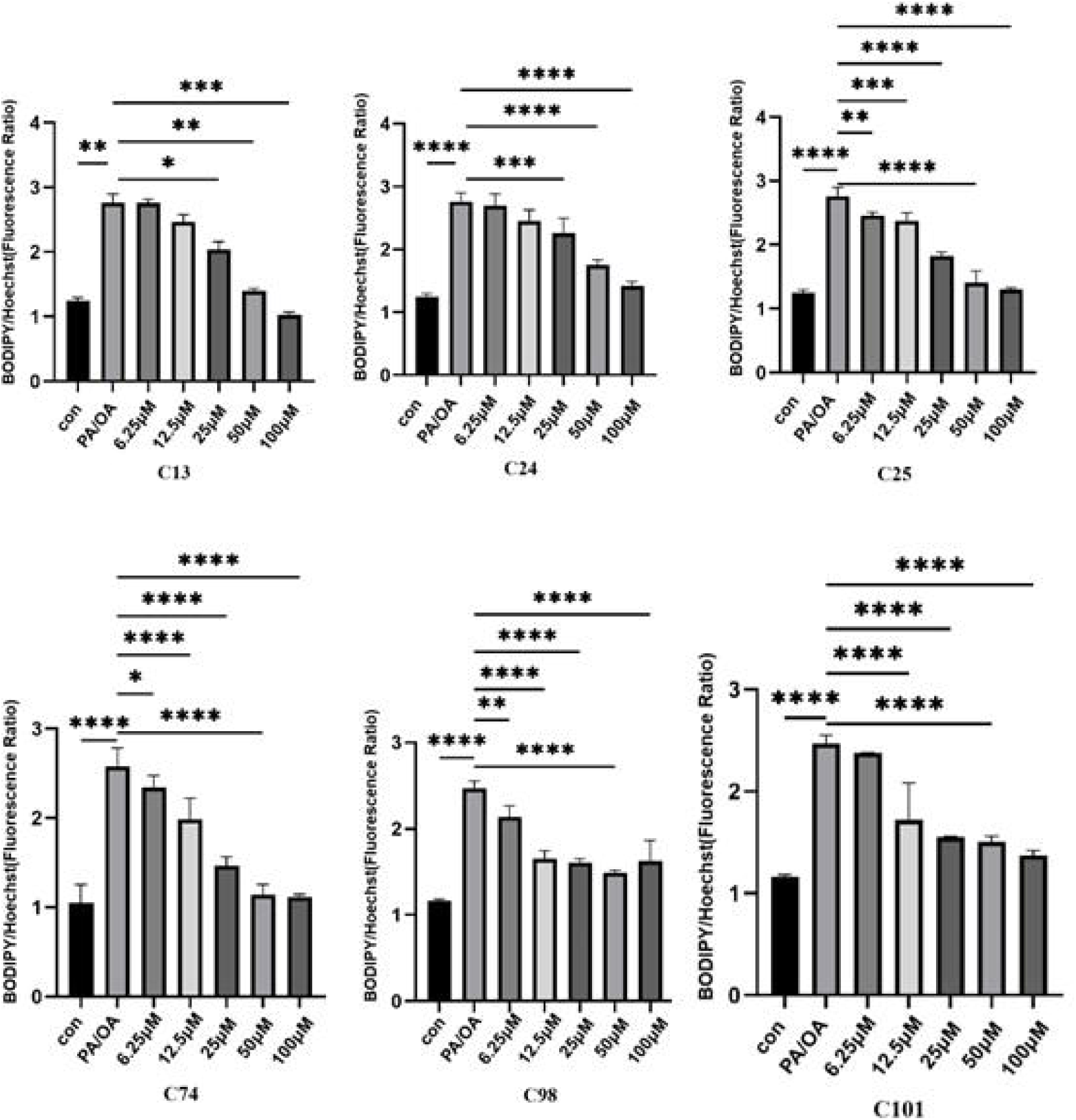
Compounds C13, C24, C25, C74, C98, and C101 significantly reduce PA/OA-induced lipids accumulation in HepG2 cells. con=vehicle alone. PA/OA = PA (100 µM) + OA (200 µM) + vehicle. Cells in compound-treatment groups are co-treated with each compound in different concentrations and PA (100 µM) + OA (200 µM). Statistics was based on One-Way ANOVA with *post hoc* Tukey comparisons between con and PA/OA group, as well as between PA/OA group and each of the compound-treatment group. (*p < 0.05, **p < 0.01, ***p < 0.001, ****p < 0.0001).

Examination of these 6 compounds by SwissADME reveals that none of the compounds complied with Lipinski’s rules since all compounds had a molecular weight greater than 500 and compounds C13, C24, C25, and C101 had a ClogP > 4.15. ^58^ Compounds C13, C24, and C25 were not considered drug-like in 2 or 3 of the remaining 4 predictors because of multiple violations to the rules and these compounds are highly hydrophobic with ClogP values ranging from 4.8-6.13. Compounds C74, C98, and C101 are drug-like in 3 out of the 4 remaining predictors (Veber, Egan, Muegge) and have lower ClogP values (4.2-4.4). ^59–61^ Interestingly, C101 is one of the most potent *C. difficile* spore germination inhibitors (IC_50_ = 0.4 ± 0.04 µM) identified to date. ^62^

### Dose- and time-dependent effect of the candidate compounds on reducing pre-accumulated lipids

With the observation of lipids accumulation-preventing effects as above, we further examined whether C13, C24, C25, C74, C98 and/or C101 had lipids-lowering effect on pre-induced lipids accumulation in HepG2 cells. As expected, all six active compounds also demonstrated both dose-(**Figure 2**) and time-dependent (**Figure S3**) lipids removal with EC_50_ values ranging between 10 and 40 µM (**Table S2**). This lipid removal effect was further visualized and confirmed with a confocal microscopy (**Figure 3**).

**Figure 2.**
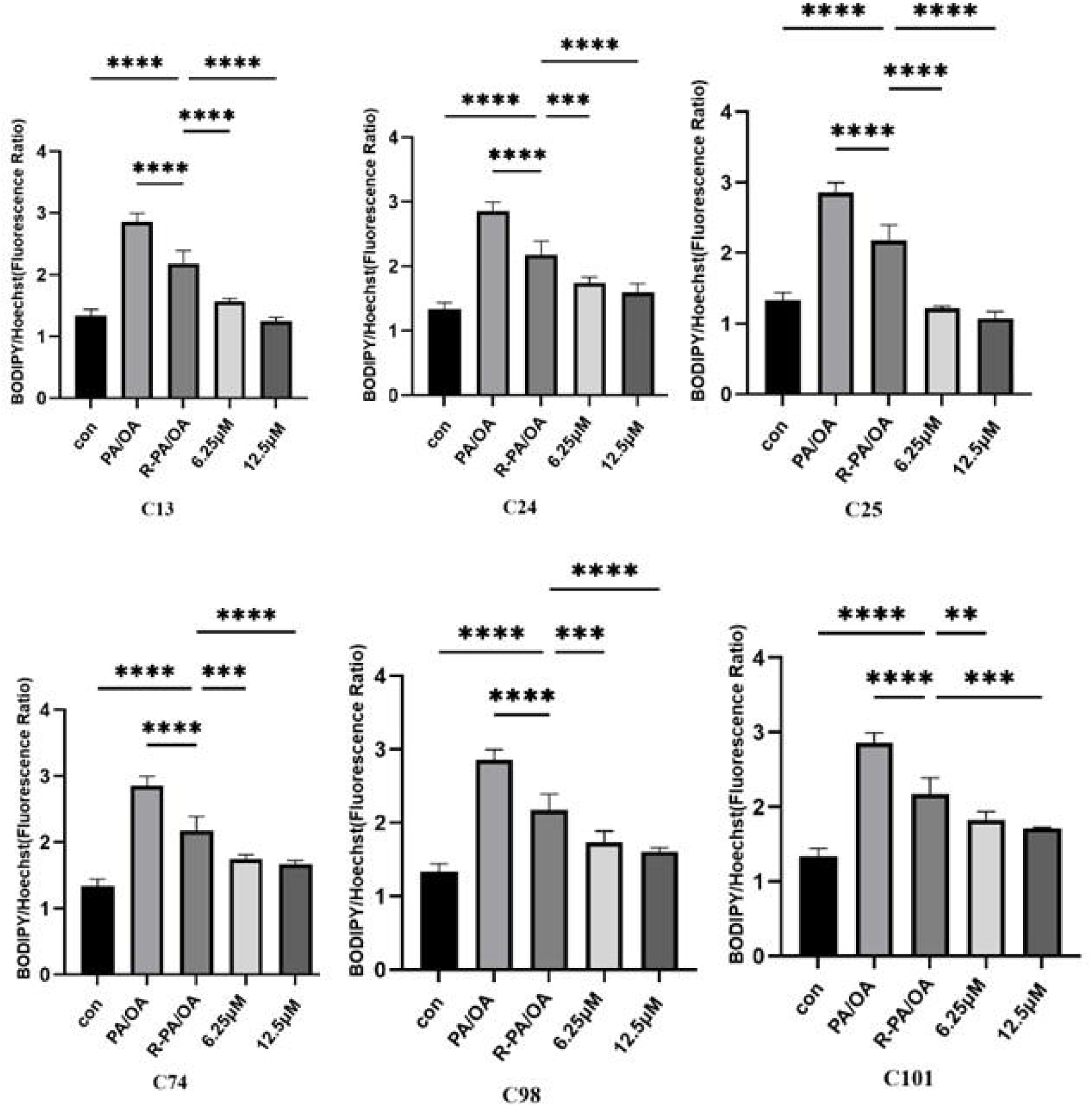
The lipid-lowering effect of compounds 13, 24, 25, 74, 98, and 101 on pre-accumulated lipids in HepG2 cells. Con = vehicle alone control for 48 hrs. PA/OA=cells were treated with PA (100 µM) + OA (200 µM) + vehicle for 48 hrs. R-PA/OA = Cells treated with PA (100µM) + OA (200 µM) + vehicle for 24 hrs, followed by a treatment with vehicle alone for additional 24 hrs. Cells in compound treatment groups are treated with PA (100µM) + OA (200 µM) + vehicle for 24 hrs, followed by a treatment with each compound in two concentrations. Statistics was based on One-Way ANOVA with *post hoc* pair-wide comparisons between con and PA/OA group, between PA/OA and R-PA/OA group, as well as between R-PA/OA and each compound-treatment group. (*p < 0.05, **p < 0.01, ***p < 0.001, ****p < 0.0001).

**Figure 3.**
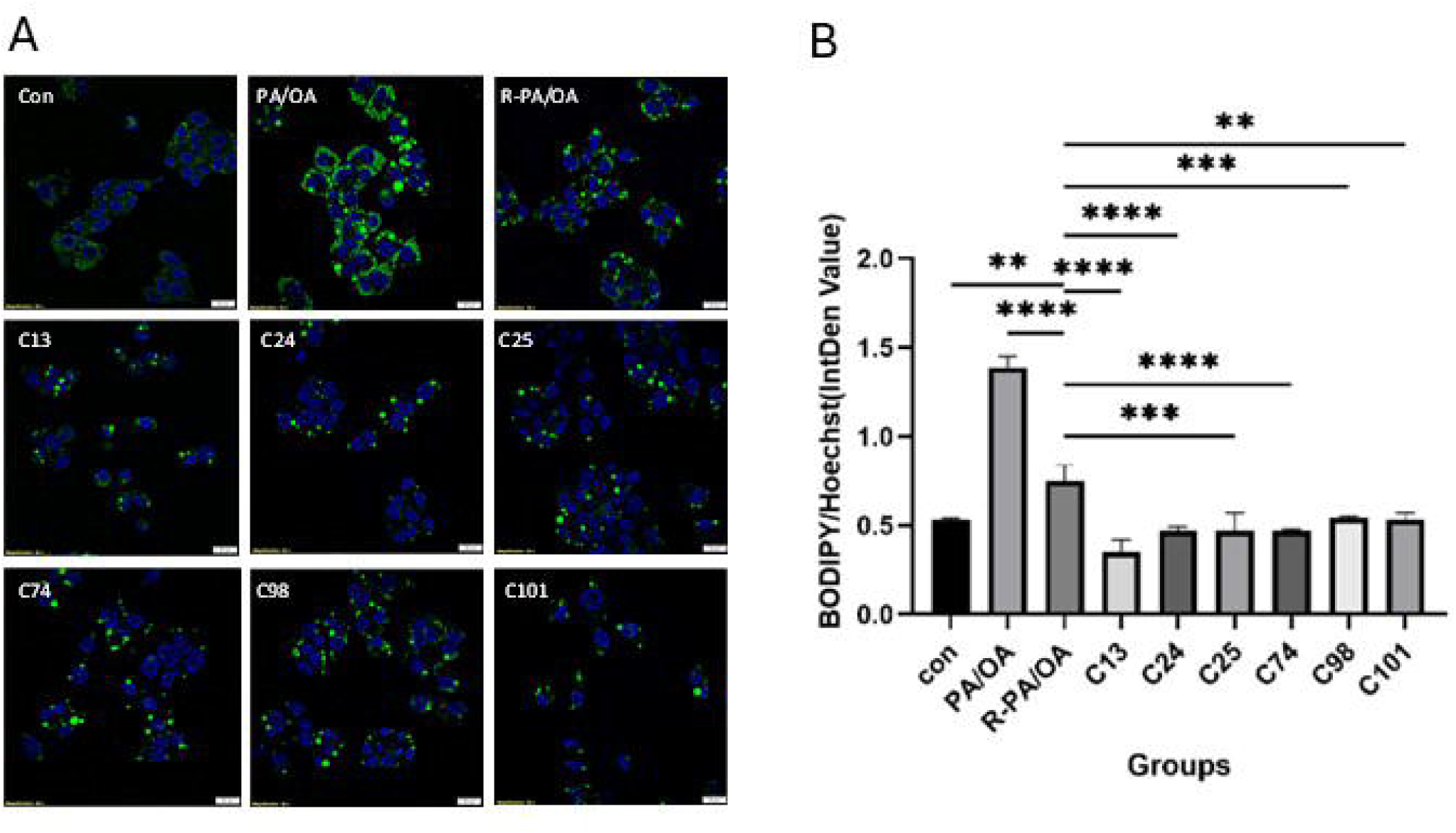
Confocal fluorescence microscopy demonstrating the BODIPY (green) and Hoechst (blue) staining (A). Data was quantified using ImageJ software (B). con = vehicle alone control for 48 hrs. PA/OA = cells were treated with PA (100 µM) + OA (200 µM) + vehicle for 48 hrs. R-PA/OA = Cells treated with PA (100 µM) + OA (200 µM) + vehicle for 24 hrs, followed by a treatment with vehicle alone for additional 24 hrs. Cells in compound treatment groups are treated with PA (100 µM) + OA (200 µM) + vehicle for 24 hrs, followed by a treatment with each compound in a fixed concentration (6.25 µM). Statistics was based on One-Way ANOVA with *post hoc* pair-wide comparisons between con and PA/OA group, between PA/OA and R-PA/OA group, as well as between R-PA/OA and each compound-treatment group. (*p < 0.05, **p < 0.01, ***p < 0.001, ****p < 0.0001). Scale bar = 20 µm.

### Triglyceride levels in cells and culture medium

To examine whether the lipid reduction effect is due to the change of intracellular TG level, and whether the reduction of intracellular TG is attributed to increased TG secretion, we measured the total TG levels in compound-treated cells and their spent media. Similar to the accumulation of neutral lipids detected by BODIPY, the intracellular TG content in the drug treatment groups were all significantly lower than the control group (**Figure S4A**). However, there was no significant difference in TG content in the medium among all groups (**Figure S4B**).

### Lipidomic analysis by thin-layer chromatography

TLC assays again showed that all six active compounds reduced intracellular TG levels significantly in HepG2, with C25 demonstrated the strongest effect (**Figure 4**). Treatments with C13, C24 and C25 also resulted in the intracellular removal of cholesterol ester. However, C101 and the structurally related compound C98, increased levels of both cholesterol ester and free cholesterol, with only C101 reaching statistical significance. As a comparison, OCA treatment only significantly decreased the intracellular TG levels, while increasing the levels of cholesterol. No statistically significant changes were observed for the accumulation of phospholipids (PL) under the conditions tested.

**Figure 4.**
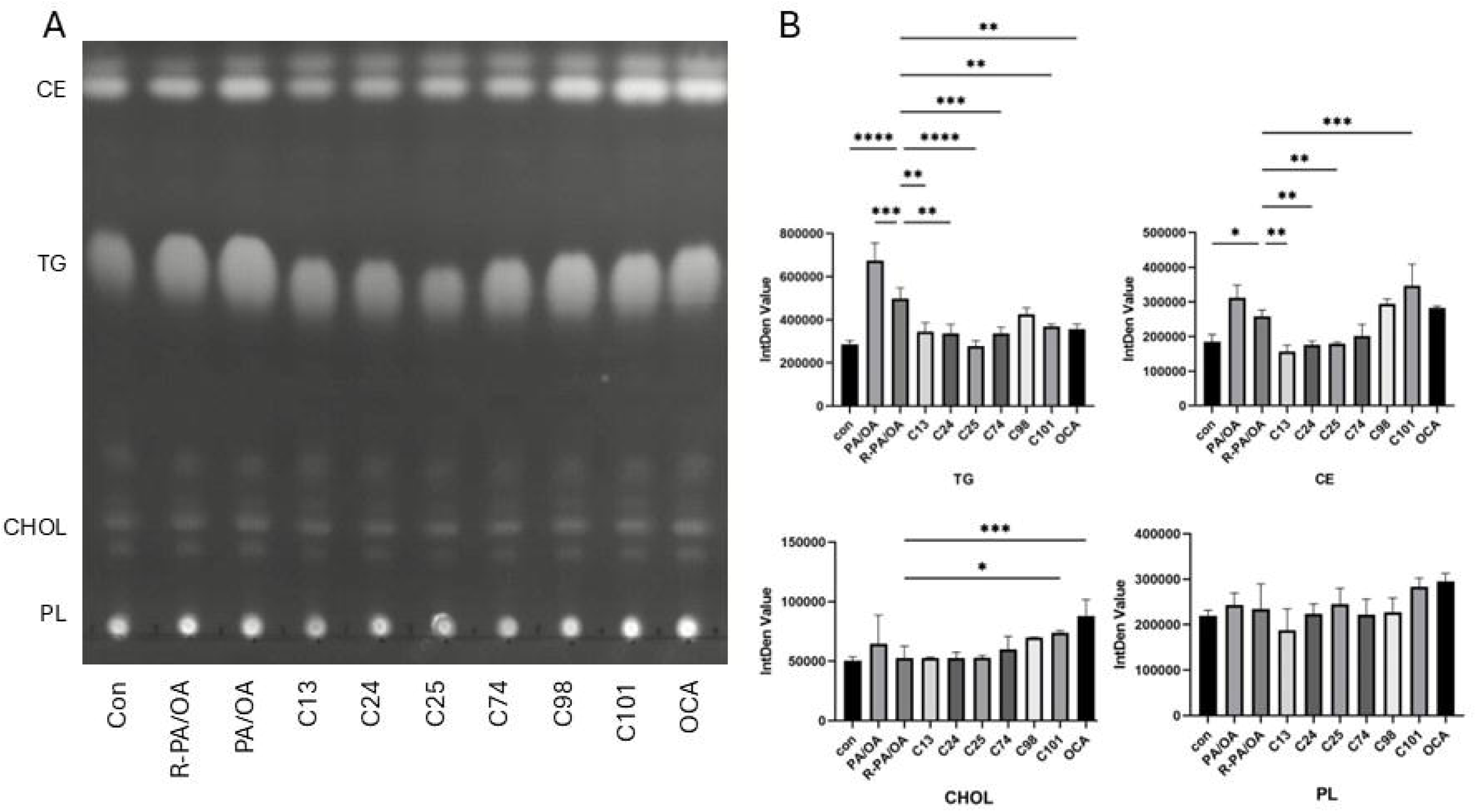
Changes of lipid fractions by compound treatment using a TLC assay. TG = triglycerides, CE = cholesterol ester, CHOL = free cholesterol, PL = phospholipids. A. TLC assay. B. Quantified data. Showing here are band intensity quantified with ImageJ. Statistics was based on One-Way ANOVA with post hoc pair-wise comparisons between con and PA/OA group, between PA/OA and R-PA/OA group, as well as between R-PA/OA and each compound-treatment (6.25 µM) group. OCA (6.25µM) was used as a control. (*p < 0.05, **p < 0.01, ***p < 0.001).

### Gene expression analysis related to lipid metabolism

To explore the potential molecular mechanism underlying the lipid-lowering effect of the six active compounds, we quantified mRNA expression level of common genes related to metabolic pathways of lipids homeostasis and metabolism (**Table S1**). Based on the transcription profile of treated HepG2 cells, the six active compounds can be separated into two clusters with C25, C24, and C74 joining together as one group, while C13, C98, and C101 cluster together with OCA in the second group (**Figure S5, Figure 5A**).

**Figure 5.**
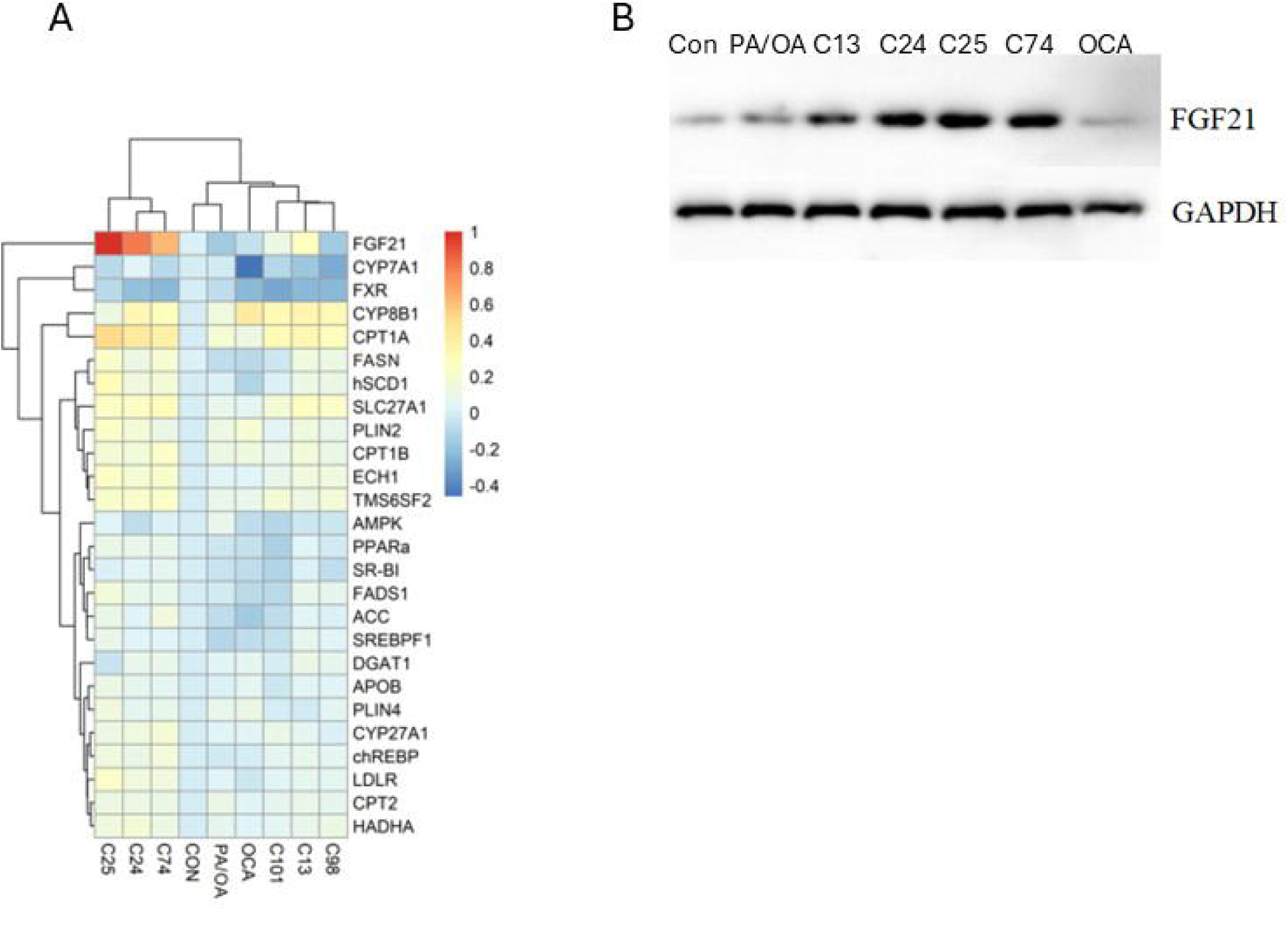
Expression pattern of mRNA (A) and protein (B) of key lipid homeostasis and metabolism genes in response to various treatments. A: Heatmap demonstrates the relative gene expression levels. B: FGF21 protein levels in each treatment group demonstrated by Western blot. con=treatment with vehicle alone for 24 hrs. PA/OA=treatment with PA (100 µM) + OA (200 µM) + vehicle for 24 hrs. Cells in compound-treatment groups are co-treated with PA (100 µM) + OA (200 µM) and each compound for 24 hrs. in a fixed concentration (6.25 µM). OCA (6.25 µM) was used as a control.

The first compound cluster was particularly characterized by a significantly increased expression of *FGF21*, while compounds in the second cluster showed a milder induction of *FGF21,* with concomitant decreases on the expression levels of *FXR* and *CYP7A1*. Western blotting also confirmed that cells treated with C24, C25 and C74 intracellularly accumulate the FGF21 hormone (**Figure 5B**).

In addition, the majority of the active compounds, but not OCA, demonstrated a significant induction of the expression of both *CPT1A* and *ECH1*, with C25 showing the strongest effect. Compounds C24, C25 and C74 also demonstrated significant expression induction for *PPAR*α, *FASN*, *hSCD1*, and *LDLR*.

### Potential effect of OCA on inhibiting C. difficile spore germination

To test the effect of OCA on *C. difficile* spores, we performed anti-germination assay *in vitro*. The assay revealed an IC_50_ of 152.0 ± 74.3 µM for *C. difficile* spore germination inhibition activity (**Figure S6**).

## Discussion

In this study, we identified six novel synthetic bile salt analogs capable of preventing and reducing lipid accumulation in hepatocytes with micromolar efficacy. Three of the compounds (C24, C25, C13) display poor drug-like properties; however, 3 compounds (C74, C98, C101) present more promising drug-like structures.

Almost all active compounds induced expression of *CPT1A* and *ECH1* which encode rate-limiting enzymes for mitochondrial lipid β-oxidation, suggesting that the lipid-lowering capacity of these compounds are associated with increased lipid metabolism. At the same time, expressions of multiple lipid synthesis genes (*e.g., FASN* and *hSCD1*) are also significantly induced by the majority of compounds, suggesting an accelerated fatty acids turnover in the cells.

Two of the active compounds (C98 and C101) behave a pattern like OCA in reducing intracellular TAG while increasing cholesterol accumulation, though C98 showed weaker activity. Notably, OCA demonstrated the strongest suppression for *CYP7A1*, confirming its activity as an FXR activator ^63^. Similarly, both C98 and C101 demonstrated a strong inhibition for *CYP7A1* level, suggesting they may share, at least in part, the same mechanism of action as OCA, which is consistent with their lipidomic profile. Whether these compounds are also FXR agonists warrants further investigation.

In contrast, C24, C25, and C74, and to a lower extent C13 seem to follow a different mechanism from OCA. These compounds are strong inducers for *FGF21*, a critical modulator for lipid metabolism. This pattern is particularly correlated with the strongest induction of both *CPT1A* and *ECH1*, target genes of FGF21 signaling, ^64–66^ suggesting that the induction of FGF21 plays the major role in reducing the lipid accumulation by these compounds. Indeed, FGF21 is a promising novel target for MASLD treatment, with a few FGF21 analogs currently being tested in various clinical trials. ^67^ The identification of robust FGF21 inducers in our study provides small molecule candidates for treating MASLD. Unfortunately, the three compounds C24, C25, and C74 are highly lipophilic and do not display drug-like properties. Further modification of these compounds is necessary to improve the pharmacological property of these compounds.

Interestingly, the well-established FXR agonist OCA, a BSA with potent activity for treating MASLD, demonstrated only a weak activity on inhibiting *C. difficile* spore germination even though OCA was able to ameliorate CDI symptoms in an obese murine model. ^45^ Contrary to OCA’s well-studied anti-MASLD activity, the mechanism behind OCA’s CDI modulatory activity is poorly understood.

Both MASLD and CDI are unmet medical needs with limited treatment options. Importantly, MASLD and CDI are clinically linked, which is believed to be mediated by the imbalanced bile acid homeostasis in the EHC and the reciprocal interactions between hepatic lipids metabolism and gut microbiota. ^49–52^ Although detailed mechanisms underlying the linkage between the two conditions have not been fully elucidated, it is an intriguing question whether reducing the risk of one disease could also prevent another. Pharmacologically, it is also unclear whether there could be a therapeutic strategy to treat or prevent both conditions simultaneously.

Different BSAs have demonstrated activities for treating MASLD/MASH and CDI prophylaxis in separate studies but no study has demonstrated any BSA with activities for preventing both diseases. Our study for the first time demonstrated that it is possible to develop novel drugs with such a dual activity. Future studies should be focused on validating the dual activity of the compound C101 *in vivo* and further improving its potency when necessary.

## Acknowledgements

This study was supported in part by the NIH grants R01AI109139, R01DK124612, and UT2AA031151.

## Abbreviations

BSA: Bile salt analogs
CD: *Clostridioides difficile*
CDCA: Chenodeoxycholic acid
CDI: *Clostridioides difficile* infection
CPT1A: Carnitine Palmitoyltransferase 1A
DMSO: Dimethyl sulfoxide
ECH1: Enoyl-CoA Hydratase 1
EHC: Enterohepatic circulation
FASN: Fatty Acid Synthase
FDA: Food and Drug Administration
FGF21: Fibroblast Growth Factor 21
FXR: Farnesoid X receptor
GAPDH: Glyceraldehyde-3-Phosphate Dehydrogenase
GLP-1: Glucagon-Like Peptide-1
HepG2: Human hepatocellular carcinoma cell line G2
hSCD1: Human Stearoyl-CoA Desaturase-1
LDLR: Low-Density Lipoprotein Receptor
MASH: Metabolic-associated steatohepatitis
MASLD: Metabolic dysfunction-associated steatotic liver disease
OCA: Obeticholic acid
OA: Oleic acid
OD580: Optical Density measured at 580 nm.
PA: Palmitic acid
PPARα: Peroxisome Proliferator-Activated Receptor Alpha
TBST: Tris-Buffered Saline with Tween 20
TG: Triglycerides
THR-β: Thyroid hormone beta receptor
TLC: Thin-Layer Chromatography
SDS-PAGE: Sodium Dodecyl Sulfate–Polyacrylamide Gel Electrophoresis.
SMILES: Simplified Molecular-Input Line-Entry System

