## Supplementary material for "Novel bile salt analogs reduce lipid accumulation in liver cells with potential to treat both metabolic dysfunction-associated steatotic liver disease and Clostridioides difficile infection": JPET Supplementary materials-4-30-26 copy.docx

**Running title:**	Bile salts prevent intracellular lipid accumulation

**Authors:**	Defeng Cai^1,2,3^, Huy Nguyen^1^, Yang Zhang^1,4^, Shiv Sharma^1^, Angel Schilke^1^, Rola Raychouni^1^, Efren Heredia^5^, Ernesto Abel-Santos^5^, Steven Firestine^1^, Wanqing Liu*^1,6^

**Affiliations:**	^1^Department of Pharmaceutical Sciences, Eugene Applebaum College of Pharmacy and Health Sciences, Wayne State University, Detroit, MI 48201, USA.

^2^Department of Clinical Laboratory (Pathology) Centre, South China Hospital, Medical School, Shenzhen University, Shenzhen, 518116, P. R. China;

^3^Institute of Biopharmaceutical and Health Engineering, Tsinghua Shenzhen International Graduate, School, Tsinghua University, Shenzhen, China.

^4^Beijing Institute of Hepatology, Beijing Youan Hospital, Capital Medical University, Beijing, China.

^5^Department of Chemistry and Biochemistry, University of Nevada - Las Vegas, Las Vegas, NV 89154, USA.

^6^Department of Pharmacology, School of Medicine, Wayne State University, Detroit, MI 48201, USA

***Corresponding author**: Wanqing Liu, PhD. Department of Pharmaceutical Sciences, Eugene Applebaum College of Pharmacy and Health Sciences, Wayne State University, Detroit, MI 48201, USA. Email: Tel:; Tel: 1-313-577-3375.

**Supplementary materials**

**Table S1.** Primers of genes related to lipid metabolism.

| Pathway | Gene | Primer Sequence（5’→3’） | Amplicon size（bp） |
| --- | --- | --- | --- |
| Lipids synthesis | ACC | F：ATGTCTGGCTTGCACCTAGTA | 106 |
|  |  | R：CCCCAAAGCGAGTAACAAATTCT | |
|  | SREBP1 | F：GCCCCTGTAACGACCACTG | 84 |
|  |  | R：CAGCGAGTCTGCCTTGATG | |
|  | ChREBP | F：AGAACCGGCGTATCACACAC | 91 |
|  |  | R：GTGCTCACGAGCCCATGAA | |
|  | FASN | F：AAGGACCTGTCTAGGTTTGATGC | 92 |
|  |  | R：TGGCTTCATAGGTGACTTCCA | |
|  | hSCD1 | F：GAGGCACCTACATTGGATGCT | 132 |
|  |  | R：CGTAGACATAGGACCGCTCA | |
|  | FADS1 | F：CTACCCCGCGCTACTTCAC | 76 |
|  |  | R：CGGTCGATCACTAGCCACC | |
| Cholesterol homeostasis | SR-BI | F：AATAAGCCCATGACCCTGAAGC | 99 |
|  |  | R：GCCCCACATGATCTCACCC | |
|  | LDLR | F：TCTGCAACATGGCTAGAGACT | 76 |
|  |  | R：TCCAAGCATTCGTTGGTCCC | |
|  | APOB | F：TGAGGAGAAGAATCGAACCCT | 89 |
|  |  | R：CTTGATTTCGTAGAGCAGACAGG | |
|  | TM6SF2 | F：GCATTGATGAGCGCCCTAATC | 83 |
|  |  | R：AGTGGGTCATAGGAGACCTCG | |
| Bile acids synthesis | FXR | F：GACTTTGGACCATGAAGACCAG | 104 |
|  |  | R：GCCCAGACGGAAGTTTCTTATT | |
|  | CYP27A1 | F：CGGCAACGGAGCTTAGAGG | 78 |
|  |  | R：GGCATAGCCTTGAACGAACAG | |
|  | CYP7A1 | F：GAGAAGGCAAACGGGTGAAC | 81 |
|  |  | R：GGATTGGCACCAAATTGCAGA | |
|  | CYP8B1 | F：CTTGTTCGGCTACACGAAGGA | 110 |
|  |  | R：GCAGGGAGTAGACAAACCTTG | |
| Lipids metabolism | AMPK | F：TTGAAACCTGAAAATGTCCTGCT | 113 |
|  |  | R：GGTGAGCCACAACTTGTTCTT | |
|  | PPARα | F：CGGTGACTTATCCTGTGGTCC | 79 |
|  |  | R：CCGCAGATTCTACATTCGATGTT | |
|  | CPT1A | F：TCCAGTTGGCTTATCGTGGTG | 98 |
|  |  | R：TCCAGAGTCCGATTGATTTTTGC | |
|  | CPT1B | F：CATGTATCGCCGTAAACTGGAC | 78 |
|  |  | R：TGGTAGGAGCACATAGGCACT | |
|  | CPT2 | F：CTGGAGCCAGAAGTGTTCCAC | 78 |
|  |  | R：AGGCACAAAGCGTATGAGTCT | |
|  | FGF21 | F：GCCTTGAAGCCGGGAGTTATT | 93 |
|  |  | R：GTGGAGCGATCCATACAGGG | |
|  | DGAT1 | F：TATTGCGGCCAATGTCTTTGC | 166 |
|  |  | R：CACTGGAGTGATAGACTCAACCA | |
|  | ECH1 | F：ATAGTGGCTTCTCGCAGACTC | 91 |
|  |  | R：CAGTGAGGCGAAGGCTAATAC | |
|  | HADHA | F：AAATTGACAGCGTATGCCATGA | 79 |
|  |  | R：GCTTTCGCACTTTTTCTTCCACT | |
|  | SLC27A1 | F：GGGGCAGTGTCTCATCTATGG | 111 |
|  |  | R：CCGATGTACTGAACCACCGT | |
|  | PLIN2 | F：ATGGCATCCGTTGCAGTTGAT | 90 |
|  |  | R：GGACATGAGGTCATACGTGGAG | |
|  | PLIN4 | F：GGAGCTGCAACCTTCGGAAA | 131 |
|  |  | R：GGACCACTCCCTTAGCCAC | |

**Table S2.** The EC50 of the lipid-reduction capacity of each compound. CI=confidence interval.

| compound | EC50（µM） | 95%CI（µM） |
| --- | --- | --- |
| C13 | **19.82** | **8.67-45.25** |
| C24 | **21.75** | **10.48-33.58** |
| C25 | **10.62** | **5.82-19.39** |
| C74 | **39.81** | **18.67-84.92** |
| C98 | **33.54** | **24.99-45.01** |
| C101 | **39.31** | **33.01-46.80** |

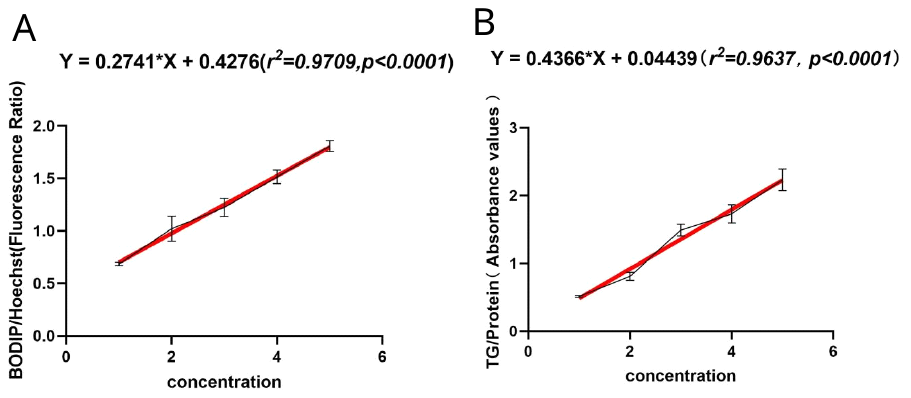
**Figure S1.** Sensitivity of the lipid quantification model in HepG2 cells. The x-axis represents the concentration of PA/OA: 1=0, 2=100µM/200µM, 3=200µM/ 400µM, 4=300µM/600µM, and 5=400µM/800µM. A) The linear relationship between PA/OA concentration and BODIPY/Hoechst fluorescence ratio. B) The linear relationship between PA/OA concentration and TG/protein ratio.

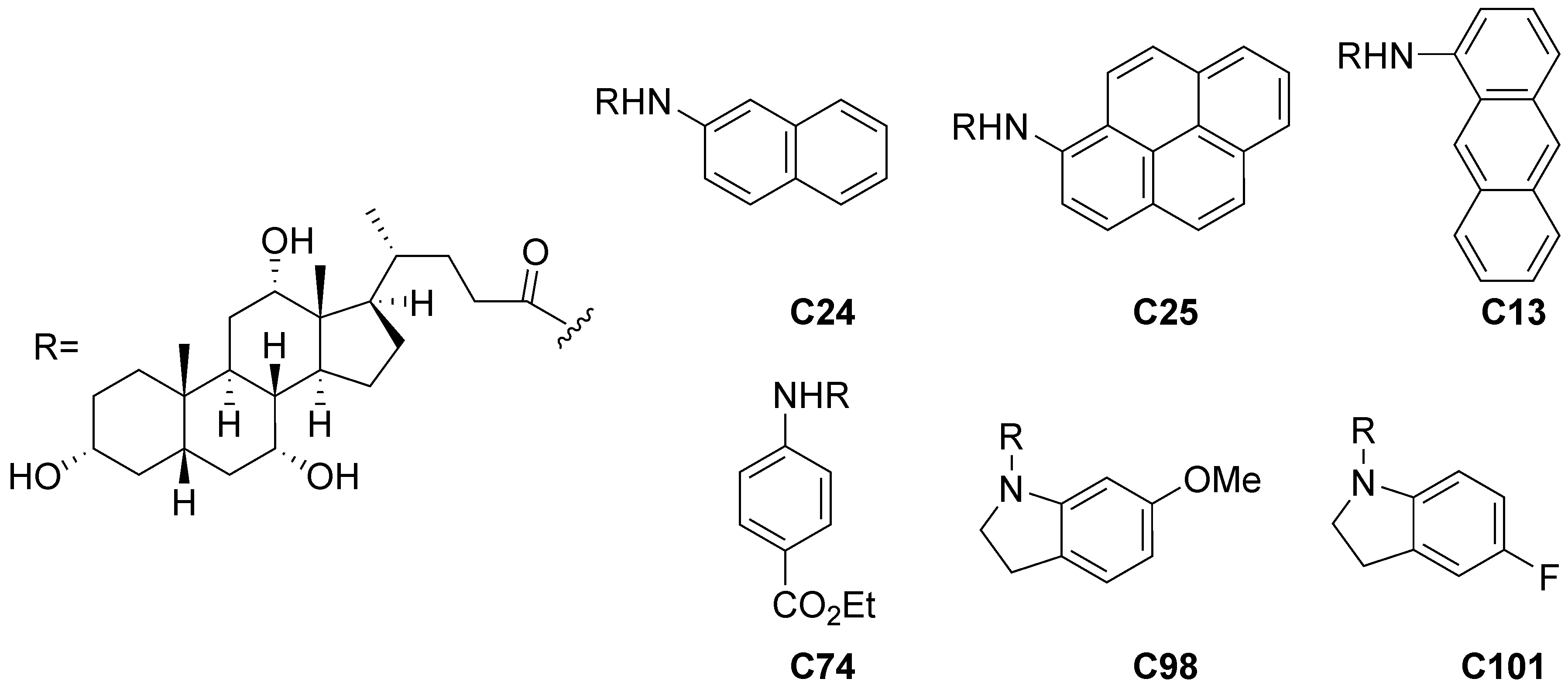

**Figure S2.** Chemical structure of representative compounds.

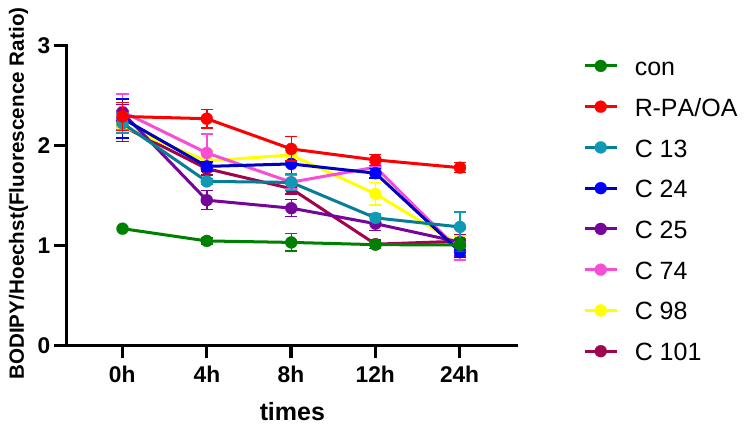

**Figure S3.** Time-dependent effect of lipid reduction of six compounds in HepG2 cells. con=vehicle alone control for 48 hrs. R-PA/OA=Cells treated with PA (100µM) + OA (200µM) +vehicle for 24hrs, followed by a treatment with vehicle alone for additional 24 hrs. Cells in compound treatment groups are treated with PA + OA + vehicle for 24 hrs, followed by a treatment with each compound (6.25 µM) in a fixed concentration for additional 4, 8, 12, or 24hrs.

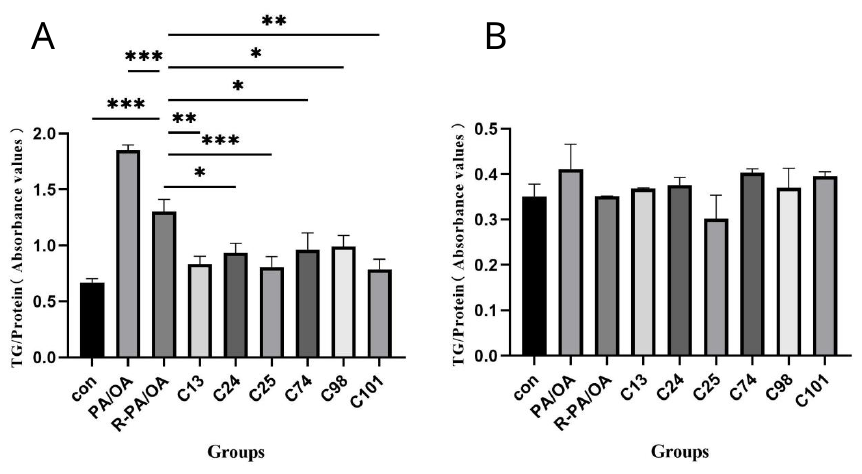

**Figure S4.** TG content in HepG2 cells (A) and the corresponding culture medium (B) of each group in the treatment model. con=vehicle alone control for 48 hrs. R-PA/OA=Cells treated with PA (100µM) + OA (200µM) +vehicle for 24hrs, followed by a treatment with vehicle alone for additional 24 hrs. Cells in compound treatment groups were treated with PA (100µM) + OA (200µM) + vehicle for 24 hrs, followed by a treatment with each compound in a fixed concentration (6.25µM) for additional 24 hrs. Statistics was based on One-Way ANOVA with a *post hoc* pair-wised Tukey test between con and PA/OA, between PA/OA and R-PA/OA, as well as between R-PA/OA and each of the drug treatment group. (*p<0.05, **p<0.01, ***p<0.001).

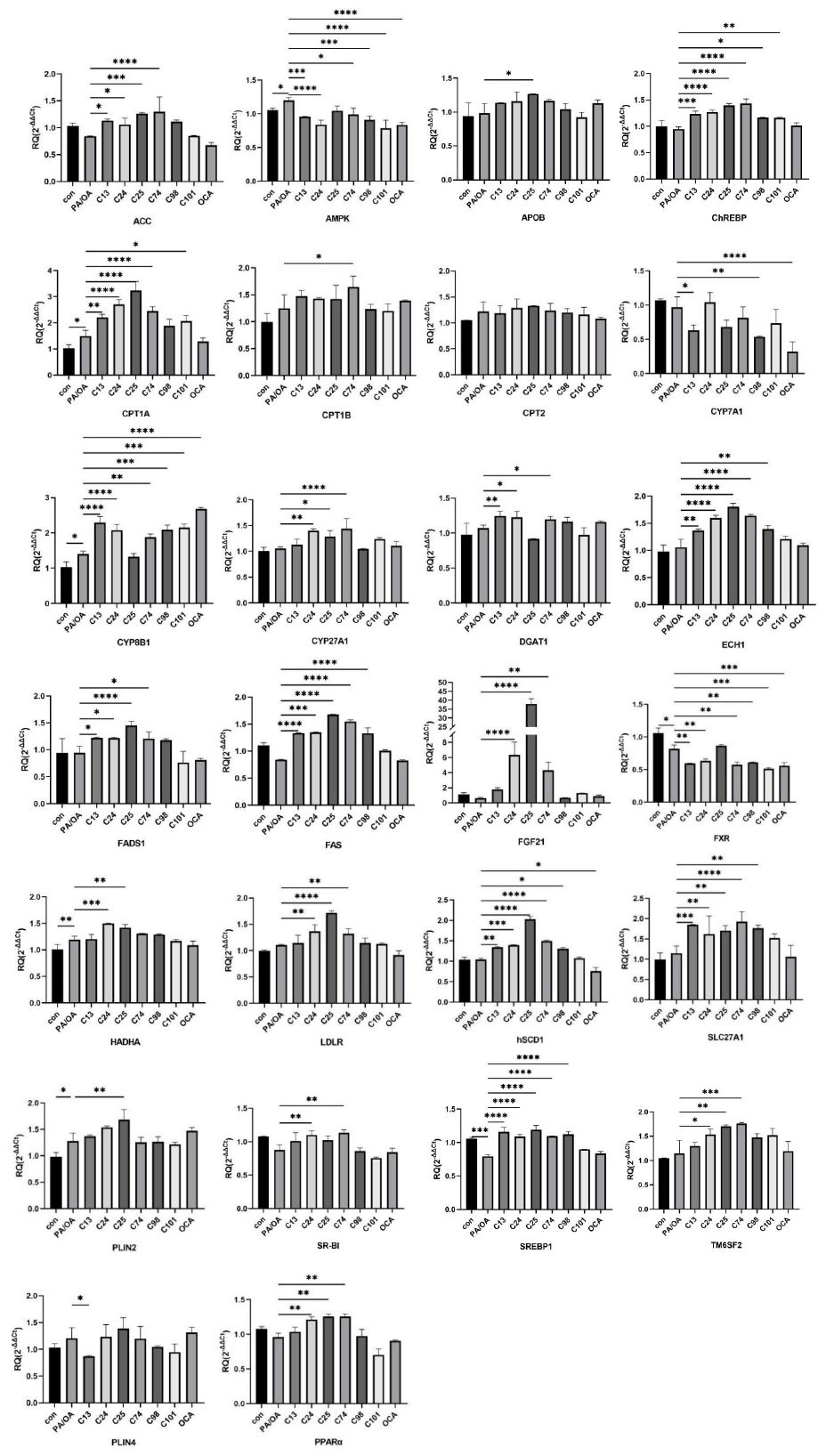

**Figure S5.** Relative mRNA expression of 26 key lipid homeostasis and metabolism genes in response to various treatments. con=treatment with vehicle alone for 24 hrs. PA/OA=treatment with PA (100µM) + OA (200µM) + vehicle for 24hrs. Cells in compound-treatment groups are co-treated with PA (100µM) + OA (200µM) and each compound for 24 hrs. in a fixed concentration (6.25µM). OCA (6.25µM) was used as a control. Statistics was based on One-Way ANOVA with a *post hoc* pair-wised Tukey test between con and PA/OA, between PA/OA and each of the drug treatment group (*p<0.05, **p<0.01, ***p<0.001, ****p<0.0001).

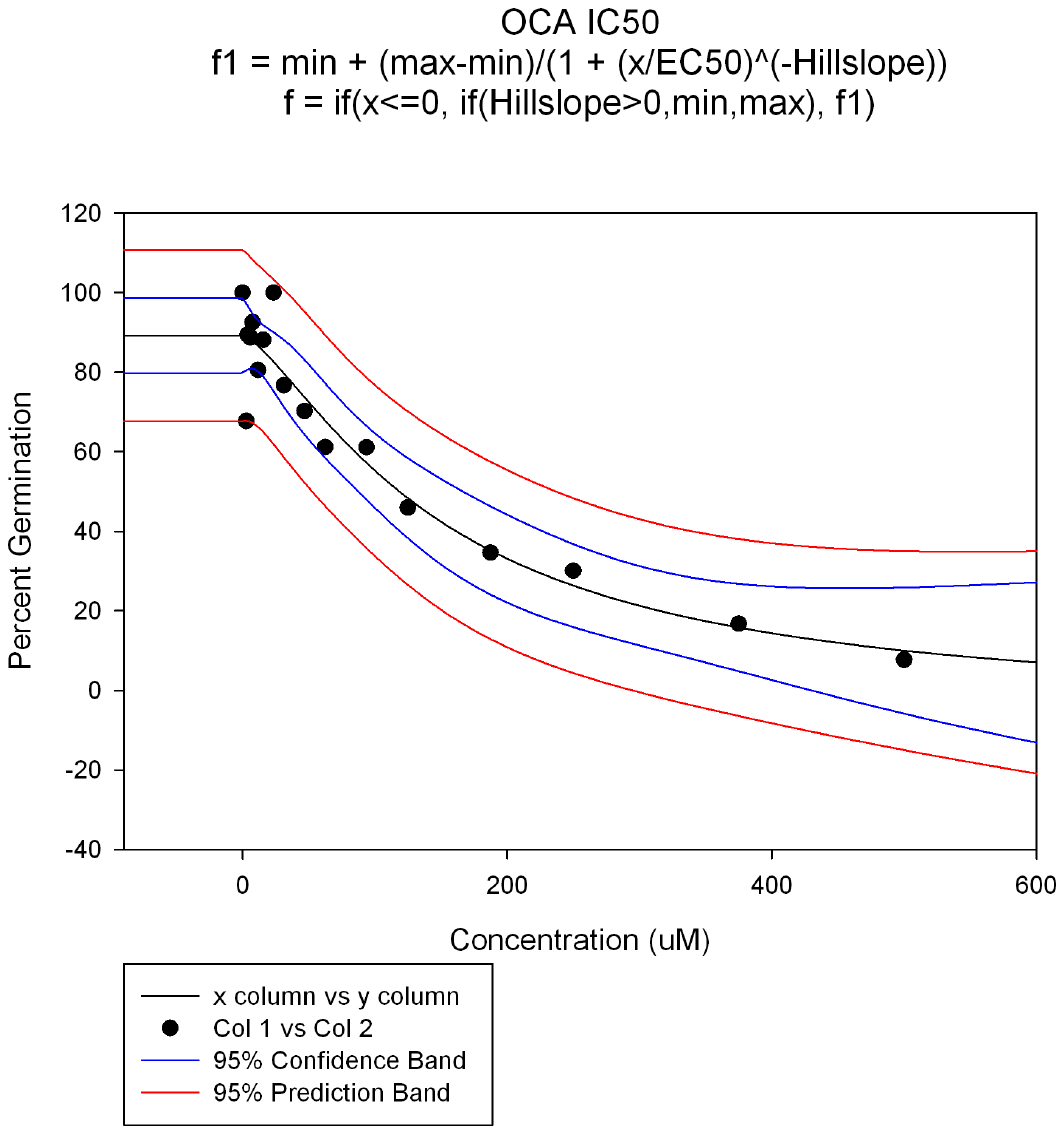

**Figure S6.** Percent germination rates of *C. difficile* spores in the presence of increasing OCA concentrations were then plotted against their corresponding OCA concentration. The data (black circles) were fitted to the four-parameter logistic function of SigmaPlot version 14 (black line). to obtain the IC_50_ value for OCA. The 95% confidence band (blue lines) and 95% prediction band (red lines) were also calculated.
